# Distribution and vulnerability of transcriptional outputs across the genome in Myc-amplified medulloblastoma cells

**DOI:** 10.1101/2021.06.07.447394

**Authors:** Rui Yang, Wenzhe Wang, Meichen Dong, Kristen Roso, Paula Greer, Xuhui Bao, Christopher J. Pirozzi, Darell D. Bigner, Hai Yan, David M. Ashley, Vasyl Zhabotynsky, Fei Zou, Yiping He

## Abstract

Myc plays a central role in tumorigenesis by orchestrating the expression of genes essential to numerous cellular processes^1-4^. While it is well established that Myc functions by binding to its target genes to regulate their transcription^5^, the distribution of the transcriptional output across the human genome in Myc-amplified cancer cells, and the susceptibility of such transcriptional outputs to therapeutic interferences remain to be fully elucidated. Here, we analyze the distribution of transcriptional outputs in Myc-amplified medulloblastoma (MB) cells by profiling nascent total RNAs within a temporal context. This profiling reveals that a major portion of transcriptional action in these cells was directed at the genes fundamental to cellular infrastructure, including rRNAs and particularly those in the mitochondrial genome (mtDNA). Notably, even when Myc protein was depleted by as much as 80%, the impact on transcriptional outputs across the genome was limited, with notable reduction mostly only in genes involved in ribosomal biosynthesis, genes residing in mtDNA or encoding mitochondria-localized proteins, and those encoding histones. In contrast to the limited direct impact of Myc depletion, we found that the global transcriptional outputs were highly dependent on the activity of Inosine Monophosphate Dehydrogenases (IMPDHs), rate limiting enzymes for de novo guanine nucleotide synthesis and whose expression in tumor cells was positively correlated with Myc expression. Blockage of IMPDHs attenuated the global transcriptional outputs with a particularly strong inhibitory effect on infrastructure genes, which was accompanied by the abrogation of MB cell’s proliferation in vitro and in vivo. Together, our findings reveal a real time action of Myc as a transcriptional factor in tumor cells, provide new insight into the pathogenic mechanism underlying Myc-driven tumorigenesis, and support IMPDHs as a therapeutic vulnerability in cancer cells empowered by a high level of Myc oncoprotein.

## Main text

*MYC* is central to human malignancies^3^, as evidenced by its wide spread gain-of-function alterations in tumor cells across two-thirds of human cancer types^6^. It has been established that oncoprotein MYC can function as a super-transcriptional factor that regulates as many as 15% of all genes in human cells, directing a transcriptional network pertinent to almost all aspects of cellular processes, including cell cycle, ribosome biogenesis, protein translation, and metabolism^2,5^. Further, over-expression of Myc oncoprotein has been shown to amplify transcriptional outputs by increasing the loading of co-activators and RNA polymerases to its target genes, leading to higher levels of transcripts from those genes^7^; however, the underlying mechanism and implication for Myc-targeting cancer therapeutic designs remain to be fully illuminated^8^. Adding another layer of complexity, recent studies have provided evidence to suggest that the far-reaching transcriptional activating and amplifying effects of Myc are only limited to certain cellular contexts^9,10^. Nevertheless, the wide spectrum of Myc’s effector genes underlie its potent oncogenic roles, and provide a strong rationale for targeting Myc for cancer therapeutics, via strategies including directly attenuating Myc’s protein abundance, modulating its regulators, and blocking its functional partners^11-15^. We speculate that delineating the transcriptional outputs across the genome in Myc-amplified tumor cells will help identify the vulnerability of Myc-driven transcriptional program and formulate a strategy for its disruption.

To address this issue, we used Myc-amplified human cancer cell lines derived from pediatric MBs (D341 and D425^16^), the most common malignant brain tumor in children and in which *Myc* amplification drives a subset of tumors (Group 3 MBs) with particularly poor prognosis^17^. We generated inducible Myc knockdown (tet-on-shMyc) cell lines such that Myc’s protein level can be tightly controlled by doxycycline in a time point- and dose-dependent manner (**Supplementary Fig. 1a**). To determine the transcriptional output and its susceptibility to a reduced level of Myc, Myc’s expression was knocked down (Myc-kd) in the cells, and nascent RNA production was fluorescence-labelled for 24 hours, followed by FACS analysis. In agreement with the finding that overexpression of Myc results in higher levels of transcripts of its targeted genes^7^, knockdown of Myc led to a measurable reduction of total RNA output in the tumor cells (**Supplementary Fig. 1b**), whereas analysis of the parental D425 cells confirmed that Dox treatment alone did not suppress nascent RNA synthesis (**Supplementary Fig. 1c**). As expected, Myc-kd led to diminished growth of MB tumor cells (**Supplementary Fig. 1d**). Notably, in D425 cells, although the reduced Myc protein level and the resultant attenuated transcriptional output were obvious five days post-induction of Myc-kd, the negative impact on tumor cell proliferation did not become apparent until day 11 (**Supplementary Fig. 1d**). This suggests that upon Myc protein’s depletion, a time lag exists between the attenuated transcriptional output and the resultant functional consequence and that the cell line was appropriate for studying the effects of Myc’s abundance on global gene transcription without other significant interference, primarily cell viability or cell cycle alterations.

To determine the global distribution and output of the gene transcription in the Myc-driven MB cells, we subjected both D425-tet-on-shMyc and D341 cell lines to induce Myc knockdown via doxycycline treatment (for eight or five days, respectively). At this time point, nascent RNAs produced were labelled with biotin for a duration of 24 hours, captured by streptavidin beads, and subjected to double-stranded cDNA synthesis and next generation sequencing (total RNA-seq) (**Supplementary Fig. 2a-b)**. To map ribosomal RNA (rRNA)-derived reads, we constructed a human ribosomal DNA (rDNA, i.e., DNA sequence that codes for ribosomal RNA) reference as previously described for a separate rRNA transcript mapping^18^, in addition to standard mappings to the whole genome and to human transcriptomic reference. Overall mapping including non-unique reads ranged from 89% to 92% in whole genome mapping with 74% to 49% of the mapped reads mapping multiple times in the control D425 and D341 cell line respectively, and a majority of those multiple mapped reads were rRNA sequences (**Supplementary table 1**).

After quantifying overall percentage of mapped reads we proceeded with analyzing only the reads mapped uniquely. We aggregated the counts to a feature level (gene level for gene-wide mapping, transcriptomic mapping for mRNA, and rDNA mapping for rRNAs). For the purpose of library depth correction, we used the counts from all three mappings together with deSeq2^19^. We fitted a negative-binomial model to the counts using deSeq2 package and corrected for multiple testing using q-value package. As expected, in the control cells, a major portion of nascent transcriptional outputs was from rDNA (**Supplementary Table 1**). When only reads uniquely mapped to the genome, but not to the rDNA sequence, were considered, transcriptional outputs varied greatly among chromosomes (**Fig. 1a, Supplementary Fig. 3a**). When the size of chromosomes were considered in this mostly diploid medulloblastoma cell line^20^, this variation remained apparent, with chr 18 and chr 19 contributing the lowest and highest transcriptional output on a per base pair basis, respectively, among the nuclear chromosomes (**Supplementary Fig. 3b, Supplementary Table 2**). Notably, the highest portion of reads (∼18%) were contributed by the mitochondrial genome (mtDNA) (**Fig. 1a, Supplementary Fig. 3a-b**). While there are hundreds of mitochondria in each cell, considering the relatively small mitochondrial genome (16,569 bp), this high level of transcriptional output from the mtDNA highlights critical roles of mitochondria in these tumor cells. Comparing the transcriptional outputs between the Myc-kd cells to the control cells revealed that Myc depletion led to various degrees of reduction in the net output of the nascent rRNA in both cell lines, in agreement with the established role of Myc on activating rRNA gene transcription^21,22^ (**Fig. 1b, Supplementary Fig. 3c**). Further analyses of the genome-wide mapped reads inform our understanding of Myc’s abundance on global transcriptional output in the following ways.

**Fig. 1.**
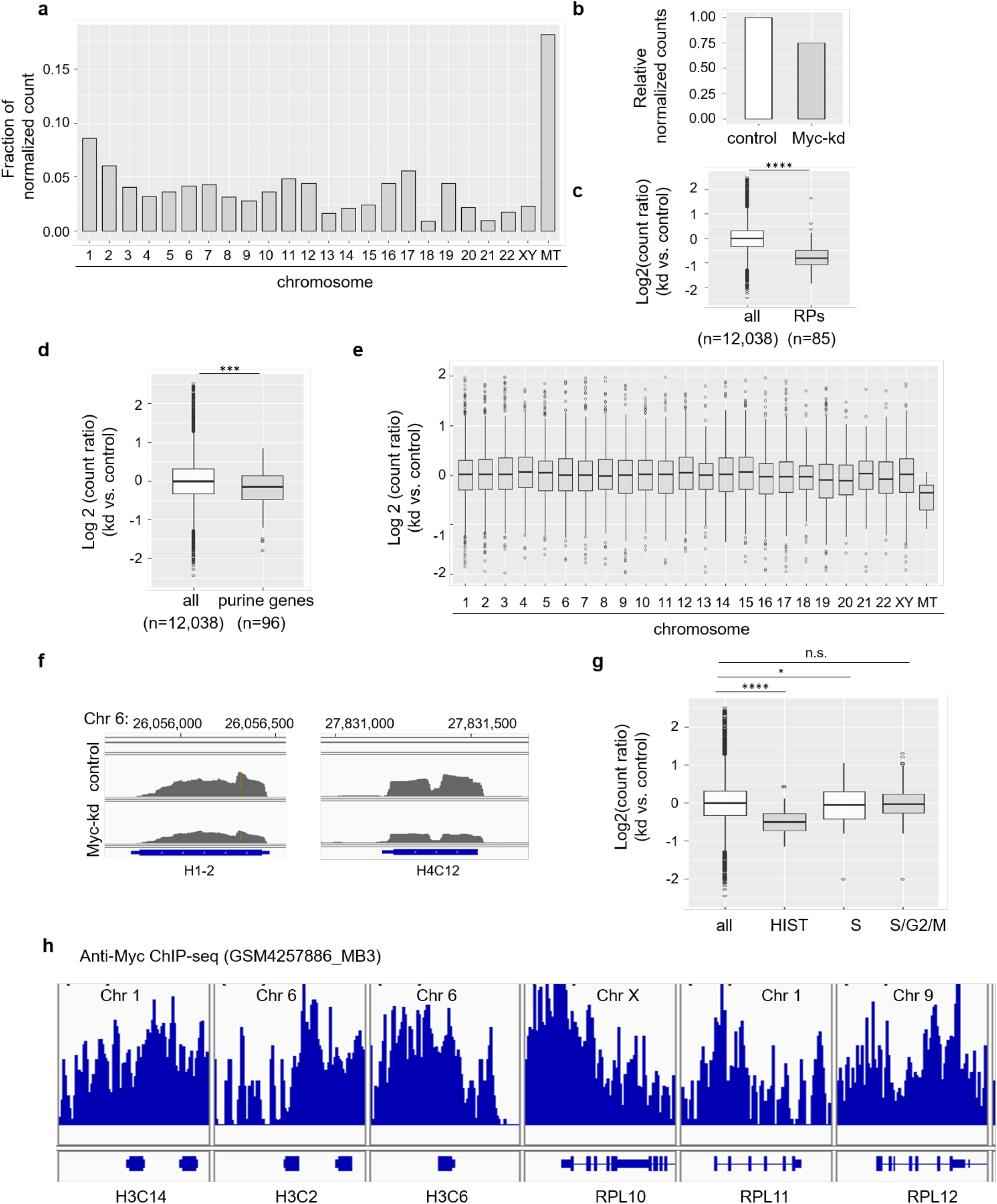
Myc depletion leads to reduced transcriptional output from genes related to ribosome biogenesis, mitochondria functionality, and histones. (**a)** The fraction of uniquely mapped reads, excluding those mapped to rDNAs, from each chromosome. (**b)** The counts mapped to the rDNA in the control versus the Myc-kd D425 cells. (**c)** The read count in the Myc-kd cells versus the count in the control cells for each gene was calculated. The ratios for all genes and for RP genes, including RPLs and RPSs, were shown. (**d)** The read count in the Myc-kd cells versus the count in the control cells for each gene was calculated. The ratios for all genes and for genes in the purine metabolism pathway were shown. (**e)** The read count in the Myc-kd cells versus the count in the control cells for each gene was calculated. The ratios for all genes, grouped by each chromosome, were shown. (**f**) Integrative Genomic Viewer (IGV) images illustrated the reduced transcript abundance for histone genes, as exemplified by H1-2 (a linker histone) and H4C12 (a core histone). (**g**) The read count in the Myc-kd cells versus the count in the control cells for each gene was calculated. The ratios for all genes and for genes encoding histones, genes that are specific for S phase, and genes that are expressed in S/G2/M phases of the cell cycle were shown. (**h**) Re-analysis of anti-Myc ChIP-seq data in Myc-driven MB tumors demonstrated the enrichment of Myc in histone genes at levels that were comparable to the enrichments in RPL genes (similar profiles were observed in three MB tumors and only images from one MB tumor are shown). ^****^p<=0.0001, ^***^p<=0.001, ^**^p<=0.01, ^*^p<=0.05.

First, in comparison to the changes in all genes, as measured by the normalized count ratio of Myc-kd vs. control, the transcriptional output from Ribosomal Protein L (RPL) and S (RPS) genes was reduced in both cell lines (**Fig. 1c**,, **Supplementary Fig. 3d**). Of note, the output from genes in the purine metabolism pathway, which are well-established transcriptional targets of Myc, by the same measurement was found to be reduced only moderately in the same 24 hour period timeframe (**Fig. 1d, Supplementary Fig. 3e**). These results, together with the above finding in rRNA transcripts, further strengthens the notion that Myc’s expression contributes to purine metabolism but is fundamentally critical to ribosome biogenesis^2^.

Second, the net output from mtDNA displayed the largest net decline among all chromosomes (**Fig. 1e, Supplementary Fig. 3f-g**), and this declined transcriptional output from the mtDNA was not simply due to the reduced read counts from the two rRNA coding genes (RNR1 and RNR2). Instead, this trend was observed in some of the 13 protein-coding genes as well as in all five tRNA genes (albeit with low read counts) (**Supplementary Table 2**). Notably, this effect of Myc’s depletion on mitochondria was not limited to genes residing in the mtDNA, as it was also observed in mitochondrial ribosomal protein genes (**Supplementary Fig. 3h**), further supporting the roles of Myc in mitochondria biogenesis^23^. We extended the analysis to nuclear genes whose protein products are exclusively localized in the mitochondria (as defined by the Human Protein Atlas), and found the overall transcriptional output from these genes were measurably reduced (as a control, the same analysis of genes encoding cytosol-specific proteins revealed no such decline) (**Supplementary Fig. 3i**). Together, these results are consistent with previous findings that Myc stimulates the expression of genes essential for mitochondrial functions and biogenesis^23,24^ in the Myc-amplified MB cells. Finally and most notably, among protein coding genes examined, genes encoding histones, including core histone H2A/B, H3 and H4, displayed notably declined transcriptional outputs (**Fig. 1f, g, Supplementary Fig. 4a, b**). Genes encoding core histones are organized as clusters in the human genome and are transcribed with a stem-loop instead of a typical poly-adenylated tail at the end^25^, making them generally undetectable by a standard profiling of typical poly-adenylated transcripts. These genes are specifically expressed in the S phase of the cell cycle and involves both transcriptional and post-transcriptional regulation^26-28^. We speculate that the attenuated transcriptional output from these histone genes was not solely due to the bystander effects of minor alterations in cell cycle, as the analysis of genes specifically transcribed in S phase or in S/G2/M phases of the cell cycle^29^ revealed no significant reduction in their abundance (**Fig. 1g, Supplementary Fig. 4b**). To further define the regulatory role of Myc in the expression of the histone genes, we analyzed previously reported ChIP-seq data in Myc-driven MBs^30^. The analysis revealed Myc’s binding to all histone genes that were detected in the nascent RNA-seq (**Fig. 1h, Supplementary Fig. 5**). Indeed, Myc binds histone genes at levels that were comparable to those in the well-established Myc-targeted genes, such as RPL genes (**Fig. 1h**). As Myc abundance has been shown to be a main predictor of Myc-regulated gene transcription^10^, these results suggest that, while it is possible that the Myc depletion-induced compromised protein translation machinery may affect the net transcriptional output from the histone genes, directly activating the transcription of histone genes is likely another key component of Myc’s oncogenic activity.

Despite the above findings, the overall limited effects inflicted by Myc’s depletion, together with the current challenges in abrogating Myc’s oncogenic function^11^, prompted us to search for an alternative target for disrupting the transcriptional output in the Myc-amplified tumor cells. We postulated that three criteria had to be met for such an alternative target: *(i)* it is strongly and positively correlated with Myc’s expression; *(ii)* it serves essential functions in Myc-overexpressing tumor cells; and *(iii)* it functions as an enzyme whose activity can be therapeutically inhibited. We utilized available gene expression profiling data from medulloblastoma (http://r2.amc.nl)^31^ to search for Myc-correlated genes/pathways. We found that within the Group 3 MBs, among the top pathways that are positively correlated with Myc are those related to RNA biogenesis/protein synthesis, and purine metabolism, in agreement with the essential roles of Myc in these cellular processes (**Supplementary Fig. 6a**). Further analysis focusing on purine metabolism found that among 36 purine synthesis enzyme coding genes, 33 of them displayed a significant (p<0.01) positive correlation with Myc (chi-square test p<0.0001) (examples in **Fig. 2a, Supplementary Fig. 6b**).

**Fig. 2.**
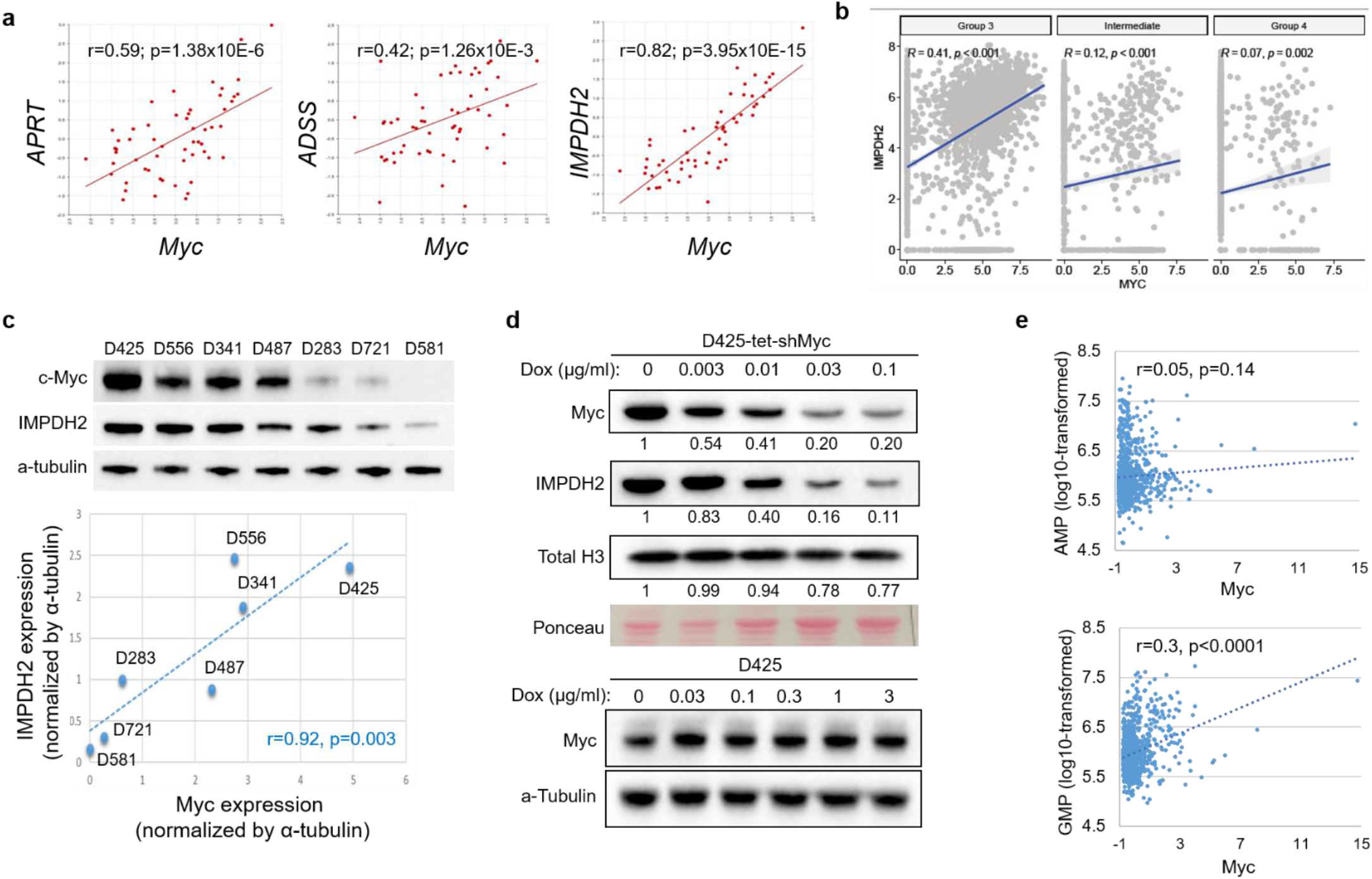
Positive correlation between Myc’s expression and the IMPDH2 expression/guanine nucleotide levels in human cancers. **(a)** Purine synthesis enzymes positively correlate with Myc expression in Group 3 MBs (n=56) (Pearson correlation co-efficient (r) and p values were shown). **(b)** Re-analysis of scRNA-seq data from MB tumors, including Group 3 (n=2138 cells), Intermediate Group (n=952 cells), and Group 4 (n=1765 cells), confirmed the positive correlation IMPDH2 and Myc expression, particularly in Group 3 tumors, at the single cell level (X and Y axis show the log2(TPM/10 + 1) normalized counts). **(c)** Myc and IMPDH2 protein levels in MB cell lines were detected by immunoblots (bottom plot shows the correlation between IMPDH2 and Myc at the protein level, with Pearson correlation r and p values). **(d)** Myc knockdown (kd) and the altered IMPDH2 expression in MB cell lines (equal amount of proteins were loaded per lane, and relative abundant of immunoblot signaling for each proteins were indicated underneath each immunoblot; note the detectable reduced levels of histone H3 in the Myc kd cells). **(e)** Positive correlation between Myc expression and the level of GMP, but not AMP, in 873 human cancer cell lines (Pearson correlation r and p values were shown).

Among purine metabolism genes, *IMPDH2* (Inosine Monophosphate Dehydrogenase 2, a rate-limiting enzyme in de novo guanine nucleotide biosynthesis), but not its homolog gene IMPDH1 - displayed the strongest correlation with *Myc* expression in MBs (**Fig. 2a, Supplementary Fig. 6c**). This positive correlation of *IMPDH2* and *Myc* were further supported by three independent lines of evidence: (i) Cells in the Group 3 MB, which are featured by high Myc expression, displayed greater *IMPDH2* expression, as revealed by the analysis of the recently reported single cell RNA sequencing (scRNA-seq) results from Group 3/4 MB patients^31,32^ (**Fig. 2b, Supplementary Fig. 6d**); and (ii) analysis of a panel of MB cells lines with known *Myc* gene status (i.e., its amplification and expression^16,33^) confirmed that the expression of these two genes are positively correlated at the protein level (**Fig. 2c**); and (iii) in Myc-knockdown in Myc-amplified MB cell lines led to correspondingly decreased levels of IMPDH2 (**Fig. 2d, Supplementary Fig. 6e**). We further examined the relationship between *IMPDH2* vs. *MYCN* or *OTX2*, signature oncogenic transcription factors in SHH Group and Group 3 MBs, respectively^31,34,35^, and found no correlation between *IMPDH2* and either gene (**Supplementary Fig. 6f**). Together, these results suggest a unique Myc-IMPDH/guanine nucleotide nexus in the Myc-amplified MB cells.

We further analyzed cancer cell line gene expression and metabolite data from the CCLE project^36,37^. These analyses revealed a similar positive correlation between *IMPDH2* and *Myc* expression in human cancer cell lines of different tissue types (**Supplementary Fig. 6g**). Most importantly, in human cell lines for which metabolic profiling data was available^37^, cellular levels of GMP, but not AMP, positively correlates with *Myc* expression (**Fig. 2e**). Together with the recently identified roles of IMPDH2/guanosine triphosphate in the pathogenesis of GBM^38^, and in a subset of lung cancers with Myc-conferred chemoresistance^39,40^, these results suggest that the IMPDH2/Guanosine triphosphate pathway is likely essential in the progression of a wide range of cancer types with high Myc expression.

GMP/GTP, in addition to being the essential building block for DNA replication, also serves as a key energy source for RNA biosynthesis^41,42^. Notably, large cell/anaplastic MBs, which are mostly associated with *Myc* amplification and poor prognosis, display prominent nucleoli (sites of rRNA/ribosome biogenesis)^43,44^, supporting a link between the aggressiveness of MBs/Myc overexpression, and robust purine metabolism / RNA biosynthesis. Furthermore, MB patients’ tumoral and cerebrospinal fluid was found to contain elevated levels of cGMP, a GTP-derived metabolic product and thus a potential indication of high intracellular GTP levels^45^. Although the histological/molecular subtypes of the MBs in this study was not determined^45^, the findings are in agreement with potentially critical roles of guanine nucleotide in at least a subset of MB tumors. Finally, unlike Myc, IMPDH2 functions as a metabolic enzyme that can be therapeutically targeted via small molecular inhibitors, such as mycophenolic acid (MPA) and Mizoribine (Miz)^46-48^. Considerable efforts have been invested in the development of IMPDH2 inhibitors, primarily for treating adverse immunity conditions, as exemplified by Miz, for treating patients of organ transplantation^46,48^. Taking advantage of these inhibitors, we utilized three MB cell lines (D341, D425, and D556) that display Myc amplification^16,33^ and high Myc expression (as shown in **Fig. 2c**) to test their response to IMPDH blockage. We note that while this study focuses on IMPDH2, these inhibitors target both IMPDHs (i.e., IMPDH1 and IMPDH2) and broadly disrupt the guanine nucleotide pathway.

As expected, treatment with MPA led to guanine nucleotide blockage (**Supplementary Fig. 7a**), and suppression of cell propagation (**Fig. 3a**). This MPA-induced suppression of MB cells can be rescued by exogenously supplied GMP/GTP (but not by its analog that can’t be converted to GMP, 8-bromo-cGMP) (**Fig. 3a**). Notably, compared to D556 and D341, D425 cells displayed lower susceptibility to MPA treatment (**Fig. 3b**). While this could be due to a variety of cell line-specific factors, the absence of functional *TP53* in the D425 cell line may have been one contributing factor ^49,50^. This possibility was supported by the finding that the exogenous expression of a dominant negative mutant of p53 (DN-p53)^51^ in the D556 cell line led to an decreased susceptibility (**Fig. 3b**), and is consistent with the role of p53 acting as a metabolite sensor of ribonucleotide shortage^52,53^. Additionally, the inhibitory effect of IMPDH blockage, but not the blockage of AMP production via an inhibitor of ADSS^54^, on MB cells was also observed in D425-derived orthotopic xenografts, as demonstrated by Miz-induced cell cycle alterations via both DAPI staining coupled with FACS analysis of cell cycle and BrdU labeling (**Fig. 3c-d, Supplementary Fig. 7b**). Finally and most notably, MB cell lines with high Myc expression demonstrated a higher susceptibility to MPA treatment when compared to those with lower Myc expression levels (**Fig. 3e**). Additionally, isogenic cell lines differing in Myc expression levels displayed Myc-correlated vulnerability to the MPA treatment (**Fig. 3f-g**), suggesting the activity of IMPDHs is uniquely important for MB cells with high levels of Myc expression.

**Fig. 3.**
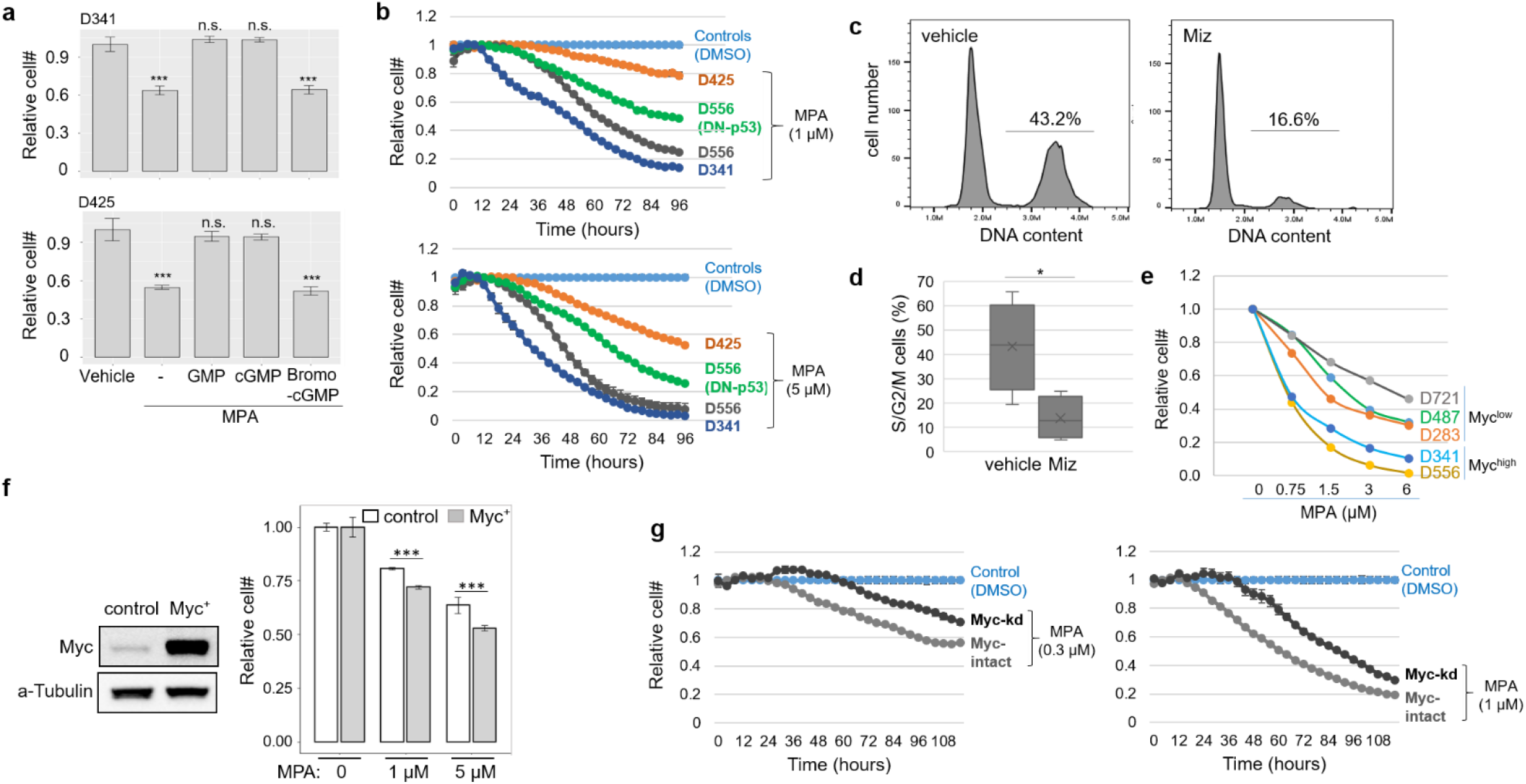
MB cell proliferation was susceptible to Inhibition of IMPDHs in vitro and in orthotopic models and this susceptibility was positively correlated with Myc’s expression. (**a)** D341 and D425 cells were treated with MPA (3 µM) with or without the presence of GMP, cGMP, or 8-brono-cGMP (10 µM each) for two days and the relative cell numbers were determined by MTA assays. **(b)** D341, D425, D556, and a derivative of D556 line overexpression a dominant-negative TP53 mutant were treated with 1 µM (top panel) or 5 µM (bottom panel) of MPA, and the cell propagation was monitored by IncuCyte (each data point represents data from triplicate wells). Note the D556-DNp53 line display a lower susceptibility to MPA compared to the D556-vector control (GFP+) cells. (**c)** D425 cell line-derived orthotopic xenografts were treated with Miz (100 mg/kg daily,, i.p. injection) for two days, and tumor cells (labelled by GFP) were harvested for Dapi staining and FACS analysis for cell cycle profiling (cell cycle profile from a representative tumor is shown for each group). (**d)** Percentages of cells in the S/G2/M phases for the vehicle- or Miz-treated group were shown (n=4 for each arm). (**e)** MB cell lines were treated with indicated doses of MPA for two days and the relative cell propagation was determined by IncuCyte (note the definition of Myc expression (high versus low) in these cell lines were based on immunoblots in Fig. 2c). (**f)** D283 cell lines with or without exogenous Myc overexpression (shown at the left panel) were treated with indicated doses of MPA for two days and cell propagation was determined by IncuCyte (data are from triplicate wells). (**g)** D341 cell lines without or with Myc-kd were treated with 0.3 µM (left panel) or with 1 µM (right panel) of MPA, and cell propagation was monitored by IncuCyte (each data point represents data from triplicate wells). ^***^p<=0.001, ^*^p<=0.05.

Collectively, these results led us to conclude that IMPDH2 met the three criteria that we initially defined in searching for an alternative target in the Myc-amplified MB cells, and prompted us to determine the effect of IMPDH2 inhibition on the transcriptional output across the genome in the Myc-amplified MB cells. We first determined the effects on MPA on the abundance of rRNAs by RT-qPCR quantification of 5S rRNA and pre-rRNA. These experiments revealed a detectable reduction of rRNAs in response to MPA (**Supplementary Fig. 8a**). We then used D425 and DN-p53-expressing D556 cell lines as the model to further determine the effects of IMPDH2 blockage on nascent RNA transcription (DN-p53 expression was introduced to minimize the complication caused by alterations in cell cycle and viability due to the high susceptibility of the parental cell line to IMPDH2 blockage). We treated the cells with MPA for 24 hours, at which point the cell proliferation phenotype was not detectable (**shown in Fig. 2b**), and measured the production of nascent RNA for a duration of 24 hours. Quantification of a marker transcript, CDKN1A, confirmed the successful capture and quantification of nascent transcripts (**Supplementary Fig. 8b**). Subsequently, total nascent RNA-seq was performed to assess the transcriptional output across the genome, following the aforementioned experimental and analysis procedures (the summary of the aligned reads in shown in **Supplementary Table 1**). The analysis revealed that in the control (vehicle-treated) cells, while the exact numbers varied, the relative transcriptional outputs across chromosomes demonstrated the same pattern in both cell lines, which was also in agreement with what was found in the control D425 cell line as shown in Fig. 1, as measured by both relative fractions and by transcriptional output on a per base pair basis (**Supplementary Fig. 8c**).

Comparing the transcriptional outputs between the MPA-treated cells versus the control cells revealed that IMPDH2 blockage leads to major reductions (reduced by ∼75% and ∼66% in D425 and D556 cell lines, respectively) in the net output of the nascent rRNAs (**Fig. 4a**). However, unlike the case of Myc depletion, the effect of IMPDH blockage on the nascent transcription of RPL and RPS genes was limited, demonstrating minimal or no reduction when compared to all genes globally (**Supplementary Fig. 9a**). These results suggest IMPDH blockage and Myc depletion can exert similarly crippling effects on ribosome biogenesis, yet with varied impacts on different components in the ribosomal machinery.

**Fig. 4.**
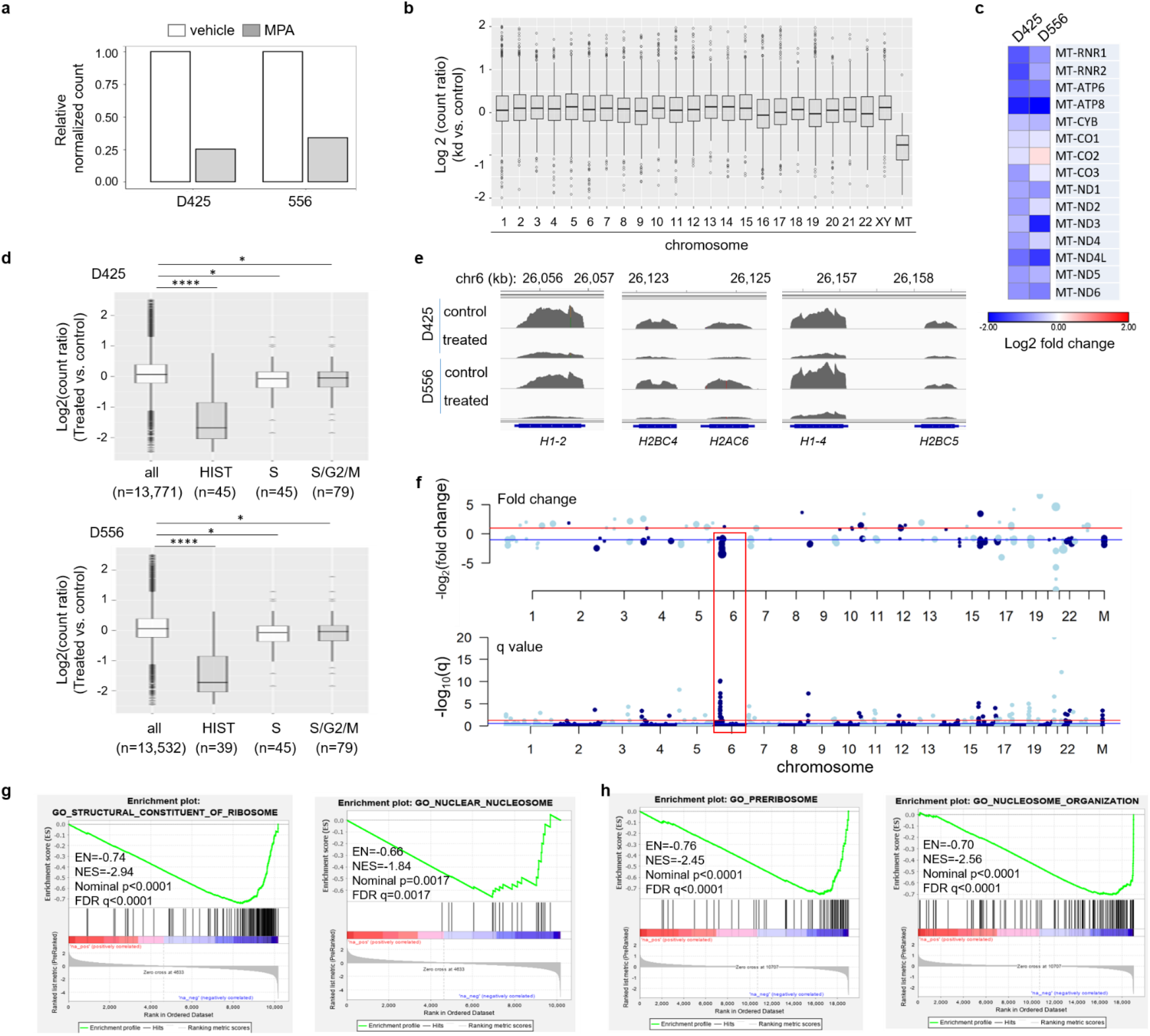
Inhibition of IMPDHs leads to altered transcriptional outputs from rDNA, mtDNA genome, and genes encoding histones. **(a)** Relative normalized counts from rDNA in the MPA-treated cells versus in the control (DMSO-treated) cells. **(b)** The read count in the Myc-kd cells versus the count in the control cells for each gene was calculated. The ratios for all genes, grouped by each chromosome, are shown (data were from both D425 and D556 cell lines). **(c)** Relative nascent transcription for genes, including two rRNA genes and 13 protein coding genes, in the mtDNA as shown by a heat map. **(d)** The read count in the MPA-treated cells versus the count in the vehicle-treated (control) cells for each gene was calculated. The ratios for all genes and for genes encoding histones, genes that are specific for S phase, and genes that are expressed in S/G2/M phases of the cell cycle are shown. **(e)** IGV images illustrate the reduced transcript abundance for histone genes, as exemplified by genes encoding linker histone genes (H1-2, H1-4) and core histones. **(f)** Manhattan plots showing the effects of MPA on nascent transcriptional output of genes across the genome (red box highlights histone gene cluster in chromosome 6). **(g)** GO pathway-based GSEA identified ribosome biogenesis and nucleosome assembly as two major pathways that were affected by Myc depletion. **(h)** GO pathway-based GSEA identified that ribosome biogenesis and nucleosome assembly were susceptible to IMPDH blockage. ^****^p<=0.0001, ^***^p<=0.001, n.s.: no significance.

Additional comparisons of the nascent RNA landscapes in the MPA-treated versus the control cells revealed IMPDH blockage differed from Myc depletion in two additional notable aspects. *First*, it exerted no measurable effects on the nascent transcription of genes involved in purine metabolism (**Supplementary Fig. 9b**), suggesting the unique importance of Myc in dictating the transcriptional output of the purine metabolism genes. *Second*, in comparison to genes across the genome, there were examples of higher transcriptional output from the promiscuous expression of intergenic regions (**Supplementary Fig. 9c)**. More noticeably, the overall transcriptional output of long intergenic non-protein coding RNAs (lncRNAs) was stimulated in response to IMPDH blockage (**Supplementary Fig. 9d-e**). The expression of lncRNA have been known to be regulated at both the transcriptional and post-transcriptional levels^55-57^. While the exact mechanism underlying and the functional significance of these stimulated transcriptional outputs remain unclear, these results suggest that, instead of a simply attenuated transcriptional output in a selected set of genes, there is a dysregulated transcriptional control across the genome in response to the blockage of IMPDHs/guanine nucleotide pathway.

When the transcriptional outputs were examined across all chromosomes, MPA treatment led to a reduction of >10% in transcriptional outputs in five chromosomes, as compared to two in the case of Myc depletion. Reminiscent of the effect of Myc depletion, the net output from mtDNA again displayed the largest net decline among all chromosomes (∼55% and 36% in D425 and D556 cell lines, respectively) (**Fig. 4b, Supplementary Fig. 10a**).

Further analysis provided two additional insights into the susceptibility of the global transcriptional output to the blockage of the IMPDHs/guanine nucleotide synthesis versus the Myc depletion. First, IMPDH blockage led to declined nascent RNA output in both RNA encoding genes (RNR1 and RNR2), and in a majority of protein coding genes in the mitochondrial genome (**Fig. 4c, Supplementary Fig. 10b**), consistent with the finding that mtDNA was the most strongly affected among all chromosomes, and highlighting the susceptibility of the transcription of mtDNA genome to the disruption of the guanine nucleotide synthesis pathway. However, unlike Myc depletion, the effect of IMPDH blockage on mtDNA’s transcriptional output was minimal for genes encoding mitochondrial ribosomal proteins (RPs) (**Supplementary Fig. 10c**), echoing the aforementioned findings in the cytoplasmic ribosomal biogenesis. In another distinction from the Myc depletion, IMPDH blockage did not affect nuclear genes encoding proteins exclusively localized in the mitochondria (**Supplementary Fig. 10d**). These results suggest IMPDH inhibition partially recapitulates effects of Myc depletion on the mitochondria, i.e., on mtDNA’s transcriptional output and mitochondrial ribosomal biogenesis. Second, MB cells treated with MPA displayed a strongly declined transcriptional output from histone genes including both core histone genes as well as linker histones and histone variants (**Fig. 4d-f, Supplementary Table 3**). A reduced output was not observed in two other categories of genes - those encoding transcriptional factors and kinases (**Supplementary Fig. 10e-f)**. More importantly, as in the case of Myc depletion, this decline in the transcriptional output from histone-encoding genes was not solely due to any subtle alteration in the cell cycle, as parallel analysis revealed no significant alterations in the transcriptional output for genes uniquely enriched in S phase or in S/G2/M phase of the cell cycle (**Fig. 4d**). Collectively, these results highlight the transcription of histone cluster genes’ susceptibility to the disruption of the guanine nucleotide synthesis pathway.

Finally, gene set enrichment analysis using sequencing reads mapped uniquely to the genome revealed that indeed, echoing the effects of Myc knockdown (**Fig. 4f**), IMPDH blockage negatively affected gene ontology pathways associated with ribosomal biogenesis as well as nucleosome packaging (**Fig. 4g, Supplementary Table 3**). Together with the aforementioned findings in various categories of genes across the genome, these results support that blockage of IMPDHs effectively leads to declined transcriptional outputs in genes primarily involved in three cellular processes: ribosomal biogenesis, mitochondrial function, and chromatin/nucleosome packaging, reminiscent of the effects of Myc depletion.

Myc plays key roles in regulating genes involved in cellular processes that are essential to tumorigenesis^1,5,7^. However, recent evidence suggests that the number of genes directly regulated by Myc at the transcriptional level, and the magnitude of Myc’s impact on transcriptional outputs likely are cellular context dependent^8-10^. We note that our study did not define a precise set of genes whose transcription is directly regulated by Myc. Instead, the findings from Myc-amplified MB cells suggest that genes whose transcriptional output is sensitive to a declined Myc abundance are primarily limited to those of fundamental cellular infrastructure. Besides the well-established roles of Myc maintaining the ribosomal machinery, Myc’s presence in these MB cells is critical in maintaining the functionality of mitochondria, as evidenced by the declined transcriptional output from the mtDNA and from nuclear genes that encode exclusively mitochondria-localized proteins essential for mitochondrial functionality. In addition, Myc’s abundance is essential in maintaining the transcription of clusters of histone genes, suggesting orchestrating the chromosome/nucleosome packaging machinery is a key aspect of Myc’s oncogenic activity. Third, despite the measurable impacts described above, the limited effects of decreased Myc abundance on the global transcriptional outputs highlights the challenge in therapeutically depleting Myc oncoprotein or blocking its transcriptional activity, and necessitates alternative targets for the purpose of anti-Myc therapies. In this regard, the susceptibility of the Myc-driven transcriptional outputs to the blockage of the IMPDHs in the MB cells nominates the IMPDHs/guanine nucleotide biosynthesis pathway as an alternative target. Further studies illuminating the mechanism underlying the global gene transcription’s vulnerability to IMPDH/guanine nucleotide pathway blockage will be critical for exploiting this metabolic pathway for treating Myc-amplified MBs or other cancers empowered by Myc overexpression.

## Materials and Methods

### Cell Lines, Plasmids, and Antibodies

MB cell lines, previously described^16,58,59^, were cultured in Improved Optimal MEM Zinc Option media (Gibco, Formula # 05-0009DJ) supplied with 10% FBS. All cell lines were maintained in a humidified atmosphere at 37°C and with 5% CO2. For inducible Myc knockdown plasmid (Sigma Mission shRNA library, clone ID TRCN0000174055) was used, and knockdown was induced by using Doxycycline (Sigma, Cat# PHR1145). The construct for exogenous Myc overexpression was previously described^60^, and the plasmid for DN-p53 was previously described^51^. For GFP expression for fluorescent label of tumor cells, a retrovirus-based expression system (MigR1, Addgene #27490) was used. Antibodies used in the study included the following: anti-Myc (Cell Signaling Technology, Cat# 5605S), anti-IMPDH2 (Abcam, Cat# ab131158), anti-H3 (Cell Signaling Technology, Cat# 9715S), and anti-α-Tubulin (Cell Signaling Technology, Cat# 3873S).

### Cell Proliferation and Cell Cycle Assays

Analyses of cell proliferation were performed using Cell Counting kit-8 (CCK-8, Cat# CK04, Dojin Laboratories, Kumamoto, Japan), or else using the Incucyte® Live-Cell Analysis (Essen BioScience, Inc., MI). For all IncuCyte-based monitoring of cell propagation, cells were labelled with GFP, and cell propagation was monitored by scanning the cell every six hours (in 96-well plate wells). Cell propagation plots were generated following the IncuCyte Bases Analysis Software provided by the manufacturer.

### Protein Extraction and Western Blots

Total cellular protein extracts were prepared using lysis buffer containing 1% SDS (w/v), 1 mM DTT, and protease inhibitor cocktail (ROCHE, Cat# 04693132001) in PBS buffer (pH7.4), plus an equal volume of 2X Laemmli loading buffer (BIO-RAD, Cat# 161-0737) supplemented with 5% 2-mercaptoethanol. Cell lysates were then heated at 100°C for 5 minutes before being used for Western blot analysis. Lysates were resolved in 12% Bis-Tris gels (SDS-PAGE, from Novex, Cat# NP0341BOX) in MOPS running buffer (Novex, Cat# NP0341BOX). Proteins were then transferred onto PVDF membranes (Immobilon, Millipore) and incubated overnight with primary antibodies at 4°C. Then blots were incubated with HRP-conjugated secondary antibody (Cell Signaling Technology, Cat# 7076, or Cat# 7074). The signaling detection was performed by enhanced chemiluminescence using the Gel Doc XR^+^ System (BIO-RAD, Hercules, USA). All immunoblot experiments shown were repeated in at least two independent experiments. The quantification was performed by using the software provided by the Gel Doc XR^+^ System.

### Mouse Xenografts and Treatments

Eight-week-old female nude mice were obtained from the Duke Cancer Center breeding core facility (derived from the Jackson Lab, stock # 007850). Mice were maintained in the Cancer Center Isolation Facility (CCIF). Briefly, 1×10^5^ cancer cells were injected into right cortex using coordinates: 2 mm posterior to lambda, 1.5 mm to the right, 2 mm deep. Five days after the injection, mice were treated with vehicle control or with Miz (Cayman, Cat# 23128) via intraperitoneally (i.p., 100 mg/kg of body weight) injection. Mice were treated twice on two consecutive days before tumors were harvested for generating single cell suspension, Dapi staining and FACS analysis for cell cycle profiling. Alternatively, mice were treated with vehicle, with Miz, or with alanosine (Medkoo Biosciences, Cat# 200130, 225 mg/kg of body weight as previously described^54^) following the same regimen; after the second treatment, mice were i.p. injected with BrdU (Cayman, Cat# 15580, 50 mg/kg of body weight) and were sacrificed 16 hours later. Tumors were harvested for anti-BrdU immunofluorescent (IF) staining using anti-BrdU antibody (Santa Cruz, Cat# sc-32323 AF594). The animal protocol was approved by the Duke University Institutional Animal Care and Use Committee (Protocol Registry Approval Number A133-19-06).

### Immunofluorescent Staining for BrdU Detection in Tumor Tissues

Immunofluorescence was performed according to an Abcam protocol (https://www.abcam.com/protocols/brdu-staining-protocol#brdu%20in%20vivo). Formalin-fixed paraffin-embedded (FFPE) section slides of tumor tissue were deparaffinized and incubated in 2M HCL for 30 minutes. After being neutralized by 0.1M sodium borate buffer pH 8.5 for 10 minutes and washed in PBS three times, sections were stained with BrdU antibody (Santa Cruz, Cat# sc-32323 AF594, 1:200) overnight and counterstained with DAPI before being mounted with mounting medium (Vectashield H-1400). Slides were imaged with a Nikon ECLIPSE TE2000-E fluorescent microscope.

### Gene Expression Correlation Analysis

MB tumors’ gene expression data used for correlation analyses, for both individual gene and for pathway correlations, was provided by the previously reported study^31^ and the analysis was performed via the R2 site: Genomics Analysis and Visualization Platform (http://r2.amc.nl). Human cancer cell lines’ gene expression data was provided by the Cancer Cell Line Encyclopedia (CCLE) project and downloaded from cBioportal^36,61,62^, and gene expression z-scores were used for expression correlation analysis. Human cancer cell line’s metabolite profiling data was from the CCLE project^37^. The MB scRNA-seq data was obtained from the previous study (GSE119926)^32^. The R package Seurat (https://www.nature.com/articles/nbt.4096) was used for single cell clustering. TPM count matrix was used to create the Seurat object. Genes expressed in at least 3 cells and cells expressing at least 200 genes were included for the analysis. “LogNormalize” with scale factor 10000 was used to normalize the filtered data. The top 2000 variable genes were selected and used for the dimension reduction steps. The first 20 PCs were used for UMAP dimension reduction.

### Nascent RNA capture, Reverse transcription and high throughput sequencing

Nascent RNA was captured with Click-iT™ Nascent RNA Capture Kit (ThermoFisher, Cat# C1-365) according to the manufactory’s protocol. Cells were labeled with EU overnight. After biotinylation of RNA and RNA capture, first strand cDNA was synthesized by Superscript IV Vilo (Thermo Cat# 11756050). Second strand cDNA was synthesized by NEB kit (Cat# E6111S). Quantitative-PCR (qPCR) analysis of the ds-cDNA was performed using CFX96TM Real-Time System (BIO-RAD, Hercules, USA), and the results were analyzed using CFX Maestro Software (BIO-RAD, Hercules, USA). Primers used for the qPCR experiments included the following: *pre-rRNA* forward: GCTCTACCTTACCTACCTGG and reverse: TGAGCCATTCGCAGTTTCAC; *5S-rRNA* forward: GGCCATACCACCCTGAACGC and reverse: CAGCACCCGGTATTCCCAGG; *CDKN1A* forward: TGTCCGTCAGAACCCATGC and reverse: AAAGTCGAAGTTCCATCGCTC; and *B2M* (as an internal control) forward: TGCCGTGTGAACCATGTG and reverse: ACCTCCATGATGCTGCTTACA. All RT-qPCR were performed in at least two independent experiments and representative results were shown. Next generation sequencing service, including library construction and sequencing, was provided by Novogene Corporation Inc. (Sacramento, CA). NovaSeq 6000 was used for PE150 sequencing for generating >=6G saw data per sample.

### Total RNA-seq Data Analysis

We mapped all the reads using Tophat v.2, allowing for 3 mismatches, read gap length 3 and read edit distance 3, versus genome reference GRCh38, RefSeq mRNA and rDNA sequences from BAC clone GL000220.1 and the primate sequences previously reported^18^. For genome wide mapping, the reads were aggregated on gene level using R/GenomicAlignments/summarizeOverlaps. We performed several different aggregations: counting only the reads falling in an exon, counting reads anywhere in a gene body, and also including 1 kb upstream to capture reads falling into promoters. This allowed us to study the consistency of exon-only counts with respect to whole-gene body count as well as producing intron and promoter counts. For transcript based mapping we simply aggregated all the counts mapped to the feature (mRNA or rRNA).

### Visualization of ChIP-seq and RNA-seq data and Gene Set Enrichment Analysis

Integrative Genomics Browser (IGV 2.8.9, released January 2020) was used for displaying and visualizing next generational sequencing data. For Myc’s binding to histone genes in MB tumors, ChIP-seq data were obtained from the recently published study (GSE143386: GSM4257878_MB1, GSM4257882_MB2 and GSM4257886_MB3)^30^. For all images illustrating the nascent RNA, the numbers of reads uniquely mapped to the genome were used for data range normalization when the IGV images were generated. Gene Set Enrichment Analysis was carrying out using GSEA version 4.1.0.^63^. Briefly, genes ranked by log2foldchange were subjected to pre-ranked analysis. Gene sets databases used included either Hallmarks (h.all.V7.3) or Gene ontology (c5.all.v7.3) and chip platform was set to MSigDB.v7.3. Pathways are considered significant if FDR<25% and nominal p value <0.05.

### Statistical Analysis

We summarized the counts at gene level for genome-wide mapping or to mRNA/rRNA for transcript based method and fit negative binomial model as described in Eqtuation 1.1 using r/deSeq2. Library normalization was done adjusting for median ratio of gene counts to geometric mean per gene. After testing for treatment effect we adjusted p-values to control for false discovery rate using r/qvalue package. Gene enrichment analysis was performed for Gene Ontology in all three domains: cellular component, molecular function and biological process using GOrilla tool, calculating hypergeometric p-values for GO terms. Enrichment was studied for mRNA transcripts significant at q-value cutoff 0.1 using all expressed mRNAs as a background dataset.

## Acknowledgement

We thank the Functional Genomics Core facility, the Duke Cancer Institute Flow Cytometry core facility, and The Cancer Center Isolation Facility (CCIF/DCI) for support to this study.

The plasmid for Myc overexpression was a kind gift from Dr. Chuan-Yuan Li, and the expression construct for the DN-p53 plasmid was a kind gift from Dr. Robert Wechsler-Reya. We thank Dr. Vidyalakshmi Chandramohan for help with the FACS analysis of cell cycle. This work was supported by The Department of Pathology, the Preston Robert Tisch Brain Tumor Center at Duke, and the National Institute of Neurological Disorders and Stroke of the National Institutes of Health, Award Number R01NS101074 (YH).

## Author Contributions Statement

R.Y, W.W., and Y.H. conceived and designed this study. R.Y., W.W., K.R., P.G., X.B., C.J.P., and Y.H. developed the methodology and performed experiments. R.Y., W.W., K.R., V.Z., and Y.H. contributed to the acquisition of data. R.Y., W.W., V.Z., Z.F., and Y.H. analyzed and interpreted the data. All authors wrote, reviewed, and/or revised the manuscript. H.Y., D.D.B., D.M.A. and Y.H. contributed to the administrative, technical, or material support. Y.H. supervised this study.

