## Supplementary material for "Distribution and vulnerability of transcriptional outputs across the genome in Myc-amplified medulloblastoma cells": Suppl Figures

**Supplementary Figures**

**Supporting Main Figure 1**

**Supplementary Figure 1. Myc knockdown (kd) in Myc-amplified MB cells. a,** Dox-induced Myc kd in D341 and D425 cell lines: 5 days after cells were exposed to the indicated doses of Dox, cell lysates were used for anti-Myc immunoblot (10 µg of lysates was loaded per lane; Ponceau staining was used to confirm comparable loading). **b,** Nascent RNA labeling and detection by FACS: Cells were exposed to Dox(100 ng/ml) for five days, at which point cells were exposed to EU for RNA labeling and one day later cells were harvested for FACS analysis. **c,** Similar treatment of parental D425 cells with Dox confirmed Dox did not attenuate the nascent RNA synthesis by itself. **d**, Myc-kd was induced by 100 ng/ml Dox, and cell propagation, as represented by relative cell numbers, was monitored by IncuCyte. Note that for D425 cells, 7 days after the experiments were started, cells were split into fresh media to allow for an extended monitoring of cell growth (each dot represents data from three wells in 96-well plates).

**Supplementary Figure 2. Nascent RNA labeling and capture for total RNA-seq. a**, Workflow for Click-iT® Nascent RNA labeling / capture for nascent total RNA-seq analysis. D425-tet-on-shMyc cells were exposed to Dox (100 ng/µL) for 7 days, at which point EU labeling was initiated for labeling nascent RNAs. One day after the labeling, cells were harvested for nascent RNA capture, ds-cDNA synthesis and total RNA-seq. **b,** RT-qPCR confirmation of enrichment of captured nascent RNAs: ds-cDNA from (a) was used for qPCR quantification of selected genes to confirm the enrichment of EU-labelled / nascent RNAs.

**Supplementary Fig. 3. The transcriptional output of mtDNA, mtDNA-located genes and genes encoding mt-localized proteins were affected by Myc depletion. a,** The fraction of uniquely mapped reads, excluding those mapped to rDNAs, from each chromosome in D341 cells. **b**, Read counts from each chromosome, excluding those mapped to rDNAs, was normalized by the size of each chromosome (counts per kb) and shown. **c,** The relative counts mapped to the rDNA in the control versus the Myc-kd D341 cells are shown. **d,** The read count in the Myc-kd cells versus the count in the control D341 cells for each gene was calculated. The ratios for all genes and for RP genes, including RPLs and RPSs, are shown. **e,** The read count in the Myc-kd cells versus the count in the control D341 cells for each gene was calculated. The ratios for all genes and for genes in the purine metabolism pathway are shown. **f**, The read count in the Myc-kd cells versus the count in the control D341 cells for each gene was calculated. The ratios for all genes, grouped by each chromosome, are shown. **g**, The ratio of the count of uniquely mapped reads (excluding those mapped to rDNAs) from each chromosome in the Myc-kd cells versus in the control cells are shown. **h**, The read count in the Myc-kd cells versus the count in the control cells for each gene was calculated. The ratios for all genes and for genes encoding mitochondria ribosomal proteins (MRPs) are shown. **i**, The read count in the Myc-kd cells versus the count in the control cells for each gene was calculated. The ratios for all genes and for genes encoding mitochondria-localized proteins (Mt) and genes encoding cytosolic proteins (Cytosol) are shown. ****$p<=0.0001$, **$p<0.01$, n.s.: $p>0.05$.

**Supplementary Fig. 4: Myc-kd led to reduced nascent transcription of histone genes. a,** GV images illustrate the distribution and abundance of sequence reads mapped to histone genes (images were from D425 cell line's data). **b,** The read count in the Myc-kd D341 cells versus the count in the D341 control cells for each gene was calculated. The ratios for all genes and for genes encoding histones (HIST, n=52), genes that are specific for S phase (n=48), and genes that are expressed in S/G2/M phases (n=35) of the cell cycle are shown. ****$p<=0.0001$, **$p<0.01$, n.s.: $p>0.05$

**Supplementary Fig. 5: Binding of Myc to histone genes in MBs.** Previously reported anti-Myc ChIP-seq data (GSM4267886_MB3) were used to generate IGV images which illustrated the binding of Myc to a panel of histone genes, including histone variant genes, core histone genes, and linker histone genes. Similar profiles were observed in three MB tumors and only images from one MB tumor (GSM4257886_MB3) are shown.

### Supporting Main Figure 2

**Supplementary Figure 6. Positive correlation between IMPDH2 and Myc expression in Group 3 MBs. a,** Pathway analysis of genes positively correlated with Myc expression in Group 3 MBs (n=56). **b,** Examples of the positive correlation between Myc expression and genes involved in the purine metabolism pathway in the Group 3 MB tumors (n=56). **c,** IMPDH1 was not positively correlated with Myc expression in Group 3 MBs. **d,** scRNA-seq analysis reveals the relatively high levels of IMPDH2 expression in cells in the Group 3 MBs (circled), in comparison to those in Intermediate and Group 4 MBs. **e,** D341-Tet-on-shMyc cells were treated with Dox for five days and immunoblots were performed (10 µg of protein lysates were loaded per lane, and the quantification of immunoblot band's intensity for each lane was labelled underneath each band). **f,** Absence of significant correlation between IMPDH2 and MYCN (SHH Group MBs, n=59) and between IMPDH2 and OTX2 (in Group 3 MBs, n=56). **g,** Positive correlation between IMPDH2 and Myc expression in human cancer cell lines:

921 cell lines for which Myc and IMPDH2 expression z-score information was available were used for analysis. For all correlation analyses, Pearson correlation r and p values are marked.

**Supporting Main Figure 3**

**Supplementary Figure 7. IMPDH blockage disrupts the guanine nucleotide synthesis pathway and affects cell cycle in orthoropic MB models. a,** MB cells were treated with 1 µM MPA for overnight and cells were harvested for metabolite measurements. The absolute concentration of each metabolite was determined, normalized by the protein concentrations, and the relative abundance of each metabolite (in the treated cell versus the vehicle-treated cells) is shown. The results confirmed IMPDH blockage-induced accumulation of its substrate (IMP) and the resultant reduction of GMP. **b,** D425-derived orthotopic xenografts were treated with Miz (100 mg/kg, via i.p. injection), and one day later another treatment of Miz at the same dose was administered while cells were also labelled with BrdU. One day later, tumor cells (GFP+) were harvested and anti-BrdU IF staining was performed. As an additional control, treatment of tumors with L-Alanosine (225 mg/kg, i.p. injection), a blocker of de novo AMP production via inhibiting ADSS, did not lead to measurable reduction in the BrdU+ cells.

**Supporting Main Figure 4**

**Supplementary Fig. 8. Validation of capture and quantification of nascent transcripts in MB cells treated with MPA. a,** RT-qPCR showing the effect of MPA (5 µM; treated for one day) on the abundances of rRNAs in D556 cells. **b,** Quantification of nascent CDKN1A transcripts in the D556 cell lines, without (left panel) or with (right panel) DN-p53 overexpression, in response to MPA treatment (5 µM, one day) confirmed the expected capture and quantification of nascent transcripts (note expected lower induction of CDKN1A nascent transcript by MPA in the D556 line expressing DN-p53). **c,** distribution of transcriptional output across the genome in the D425 and D556 cell lines.

**Supplementary Fig. 9 IMPDH blockage leads to increased transcriptional output in selected intergenic regions and in intergenic LncRNAs. a,** The read count in the MPA-treated cells versus the count in the vehicle-treated (control) cells for each gene was calculated. The ratios for all genes and for genes encoding ribosomal proteins, including RPLs and RPSs, are shown. **b,** Similar analysis to those in (a) for the genes in the purine pathway. **c,** IGV images illustrate the examples of MPA-induced transcription in intergenic regions. **d,** Similar analysis to those in (a) for detectable lncRNA-encoding genes. **e,** IGV images illustrated the examples of MPA-induced transcription in intergenic lncRNAs. ****$p<=0.0001$, *$p<=0.05$, n.s.: no significance.

**Supplementary Fig. 10. IMPDH blockage leads to attenuated transcriptional output from mtDNA and from gene encoding histones. a**, Alterations in transcriptional outputs from all chromosomes in D425 and D556 cells in response to MPA. **b,** IGV images illustrated the MPA-induced reduction in the transcriptional output from the mtDNA. **c,** The read count in the MPA-treated cells versus the count in the vehicle-treated (control) cells for each gene was calculated. The ratios for all genes and for genes encoding mitochondrial Ribosomal proteins, including RPLs and RPSs, are shown. **d,** Similar analysis to those in (c) for the genes that encode proteins that are uniquely mitochondria-localized. **e,** IGV images illustrated the MPA-induced reduction in the transcriptional output from histone-encoding genes. n.s.: no significance.

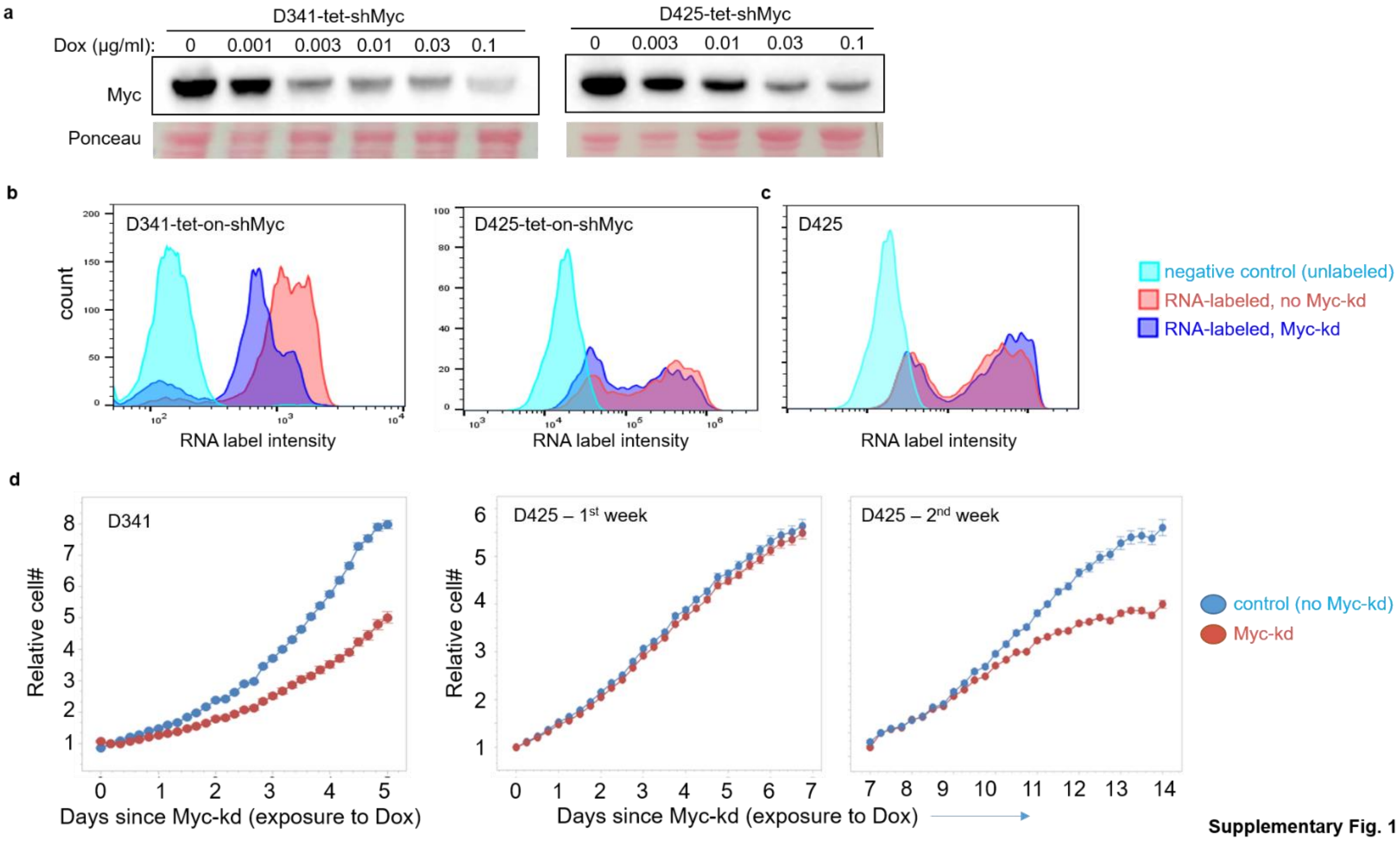


**Supplementary Fig. 1**

**a**

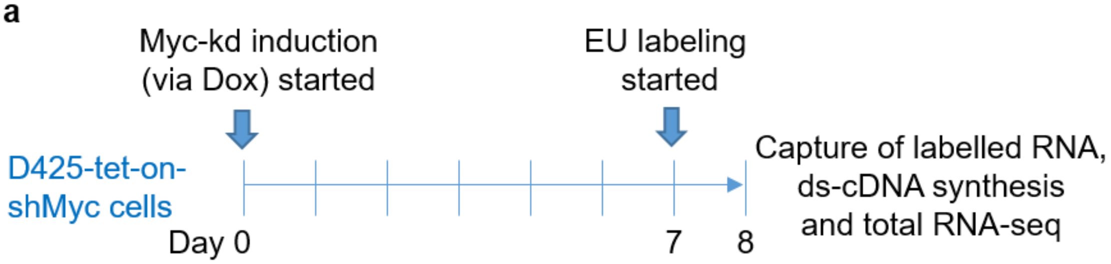


**b**

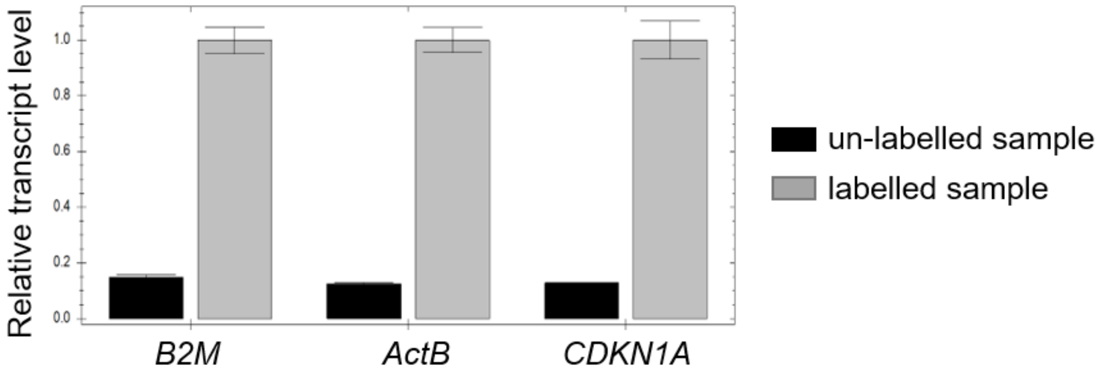


**Supplementary Fig. 2**

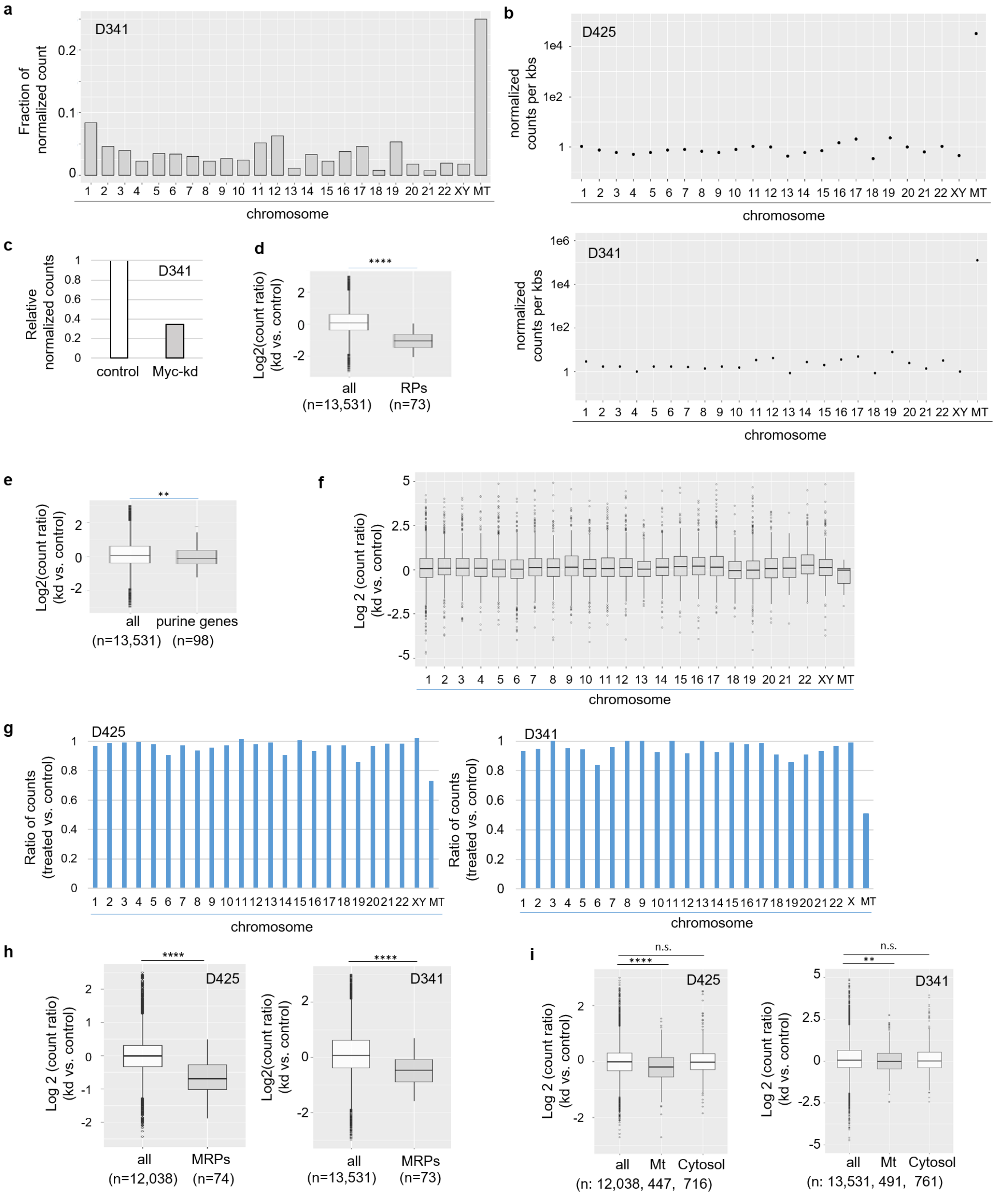


**Supplementary Fig. 3**

a

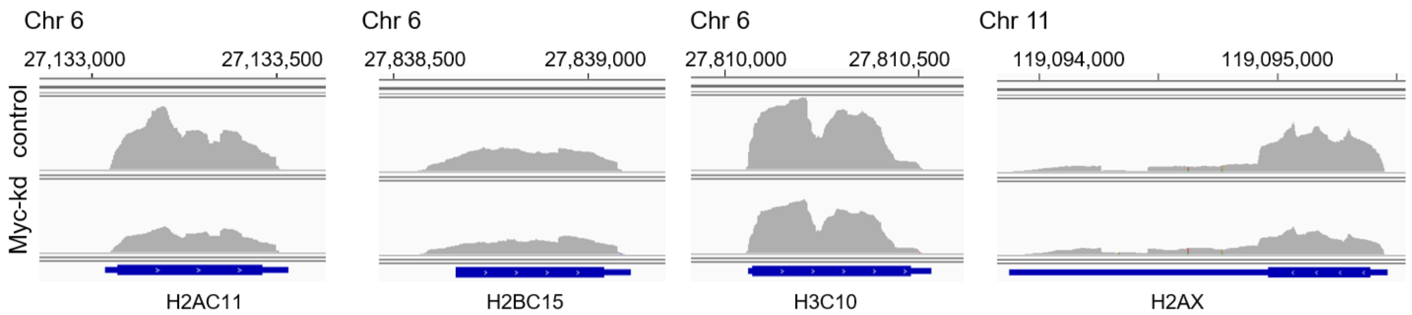


b

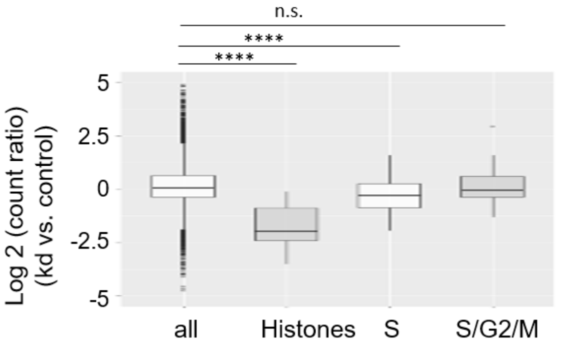


**Supplementary Fig. 4**

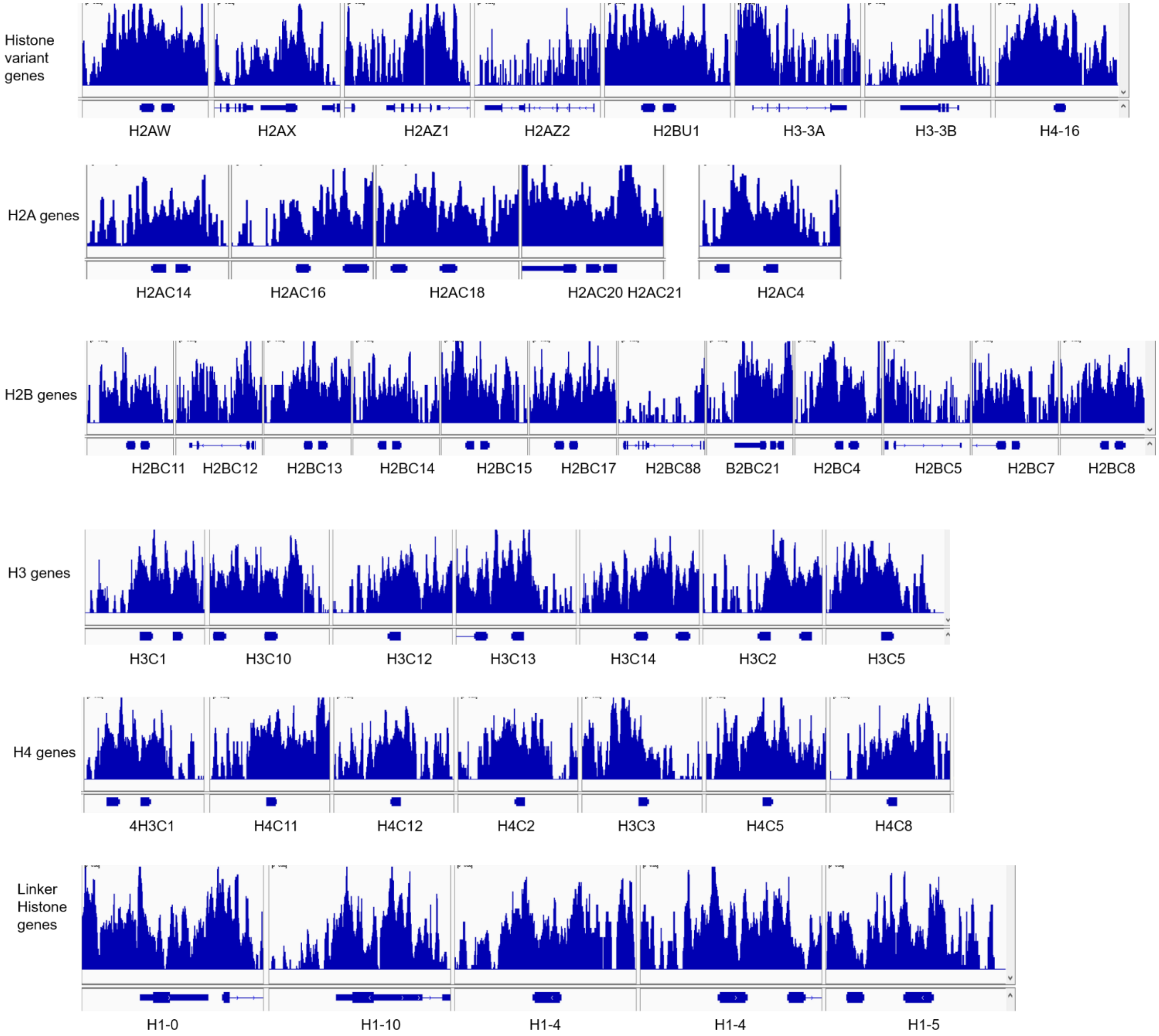


**Supplementary Fig. 5**

**a**

| series# | KEGG geneset | p value |
| --- | --- | --- |
| 1 | Ribosome | 8.50E-116 |
| 2 | Ribosome_biogenesis_in_eukaryotes | 2.50E-19 |
| 3 | RNA_transport | 6.00E-13 |
| 4 | RNA_degradation | 1.80E-09 |
| 5 | Aminoacyl_tRNA_biosynthesis | 5.50E-07 |
| 6 | Purine_metabolism | 1.10E-05 |
| 7 | Huntington_s_disease | 1.40E-05 |
| 8 | Spliceosome | 1.90E-05 |
| 9 | RNA_polymerase | 5.20E-05 |
| 10 | Parkinson_s_disease | 7.80E-05 |

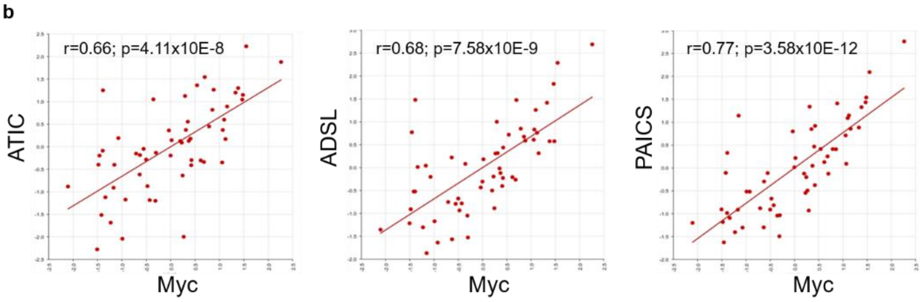


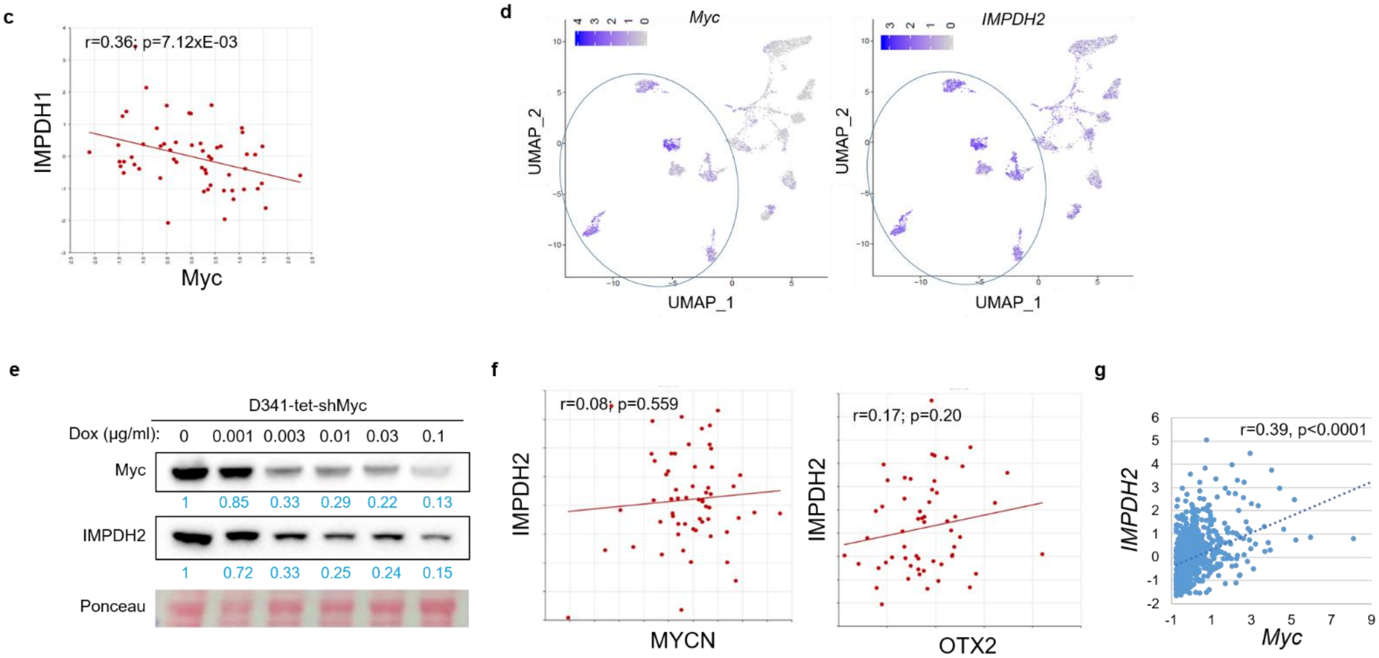


**Supplementary Fig. 6**

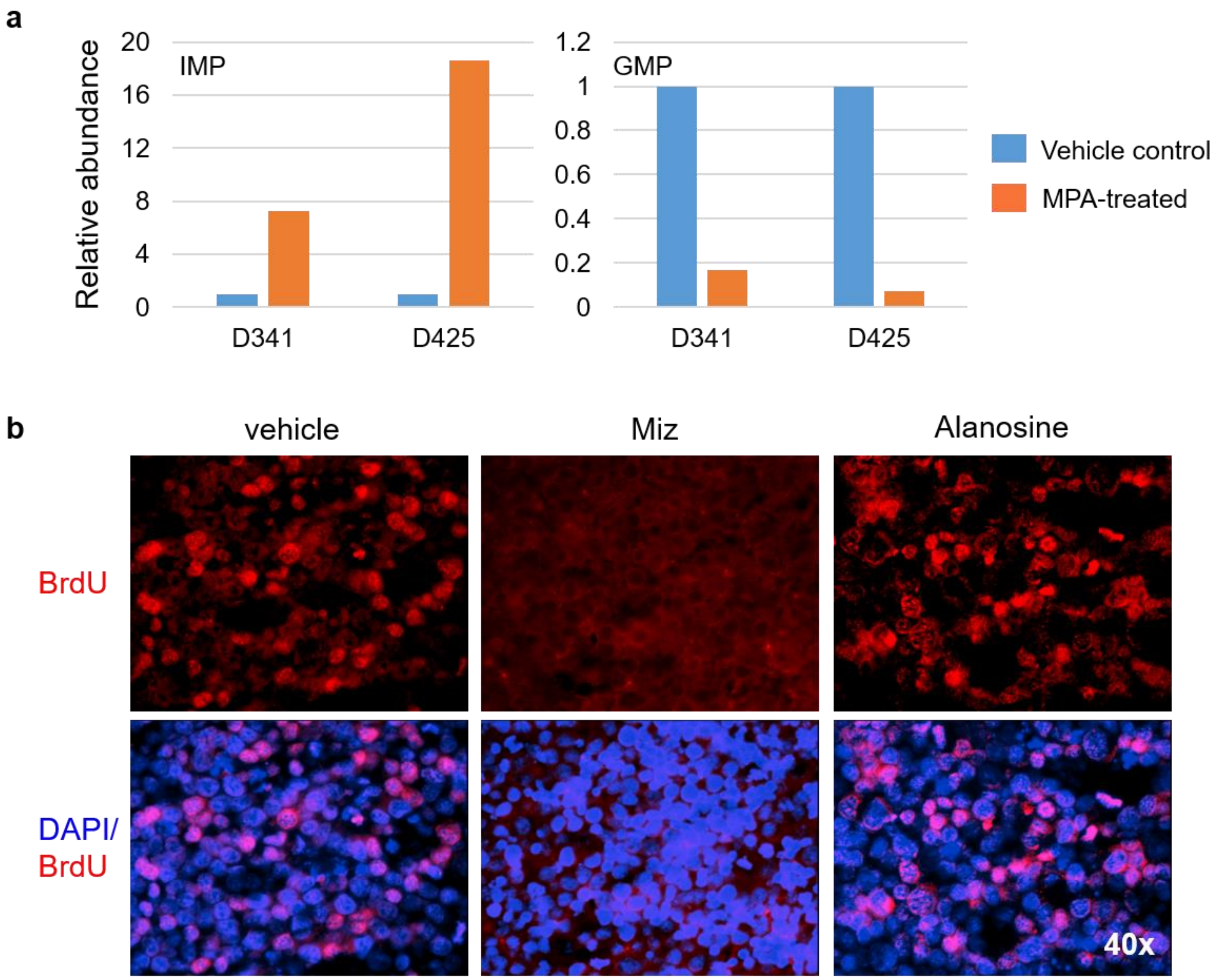


**Supplementary Fig. 7**

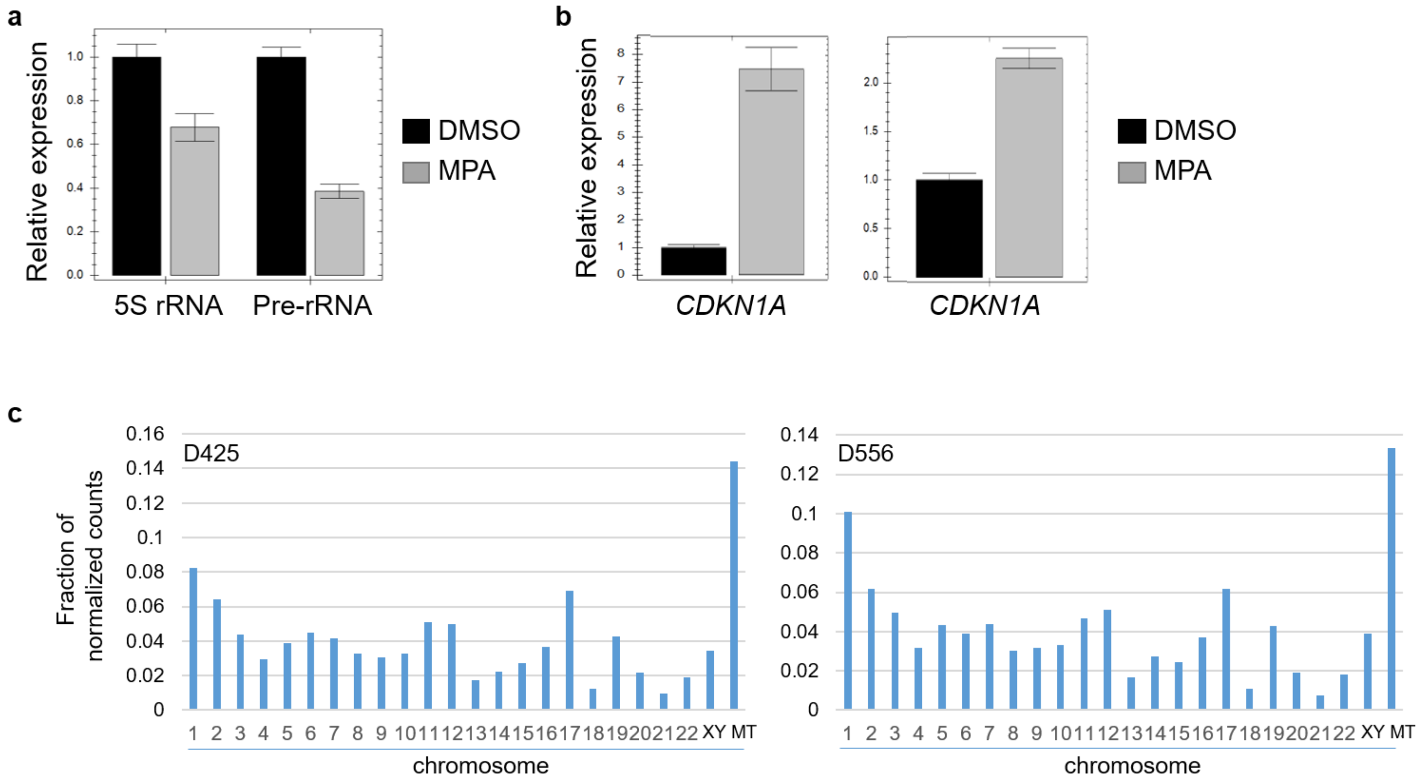


**Supplementary Fig. 8**

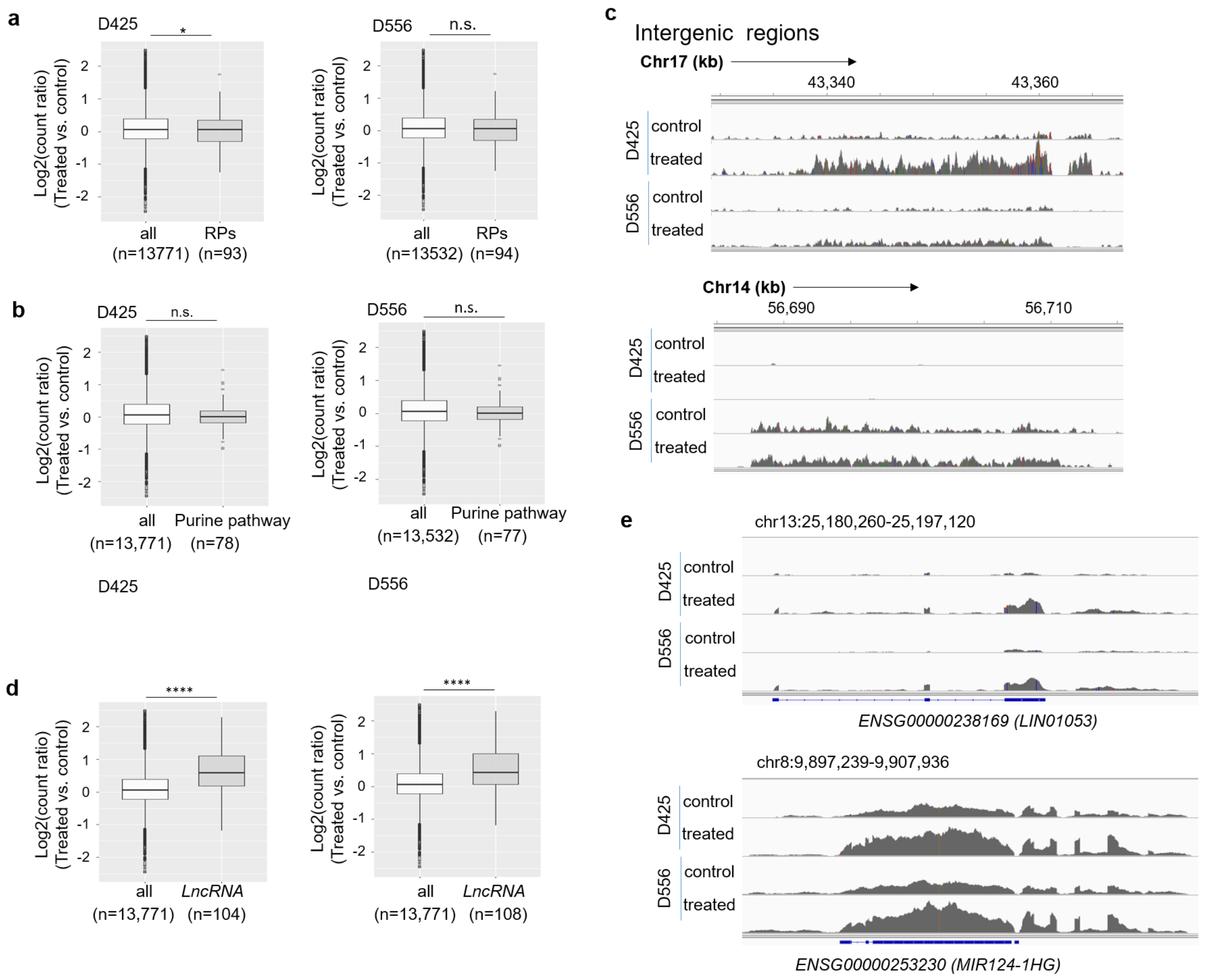


**Supplementary Fig. 9**

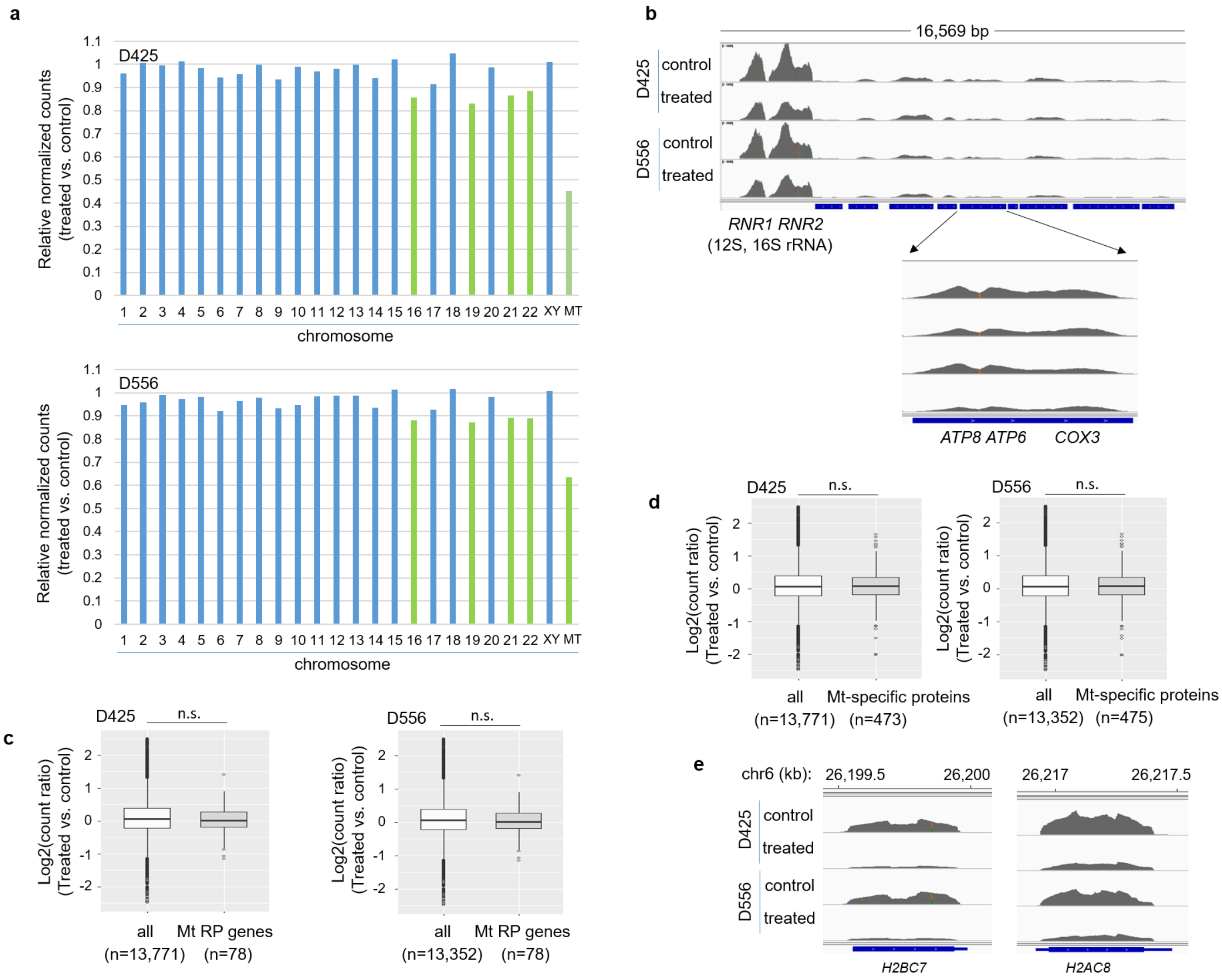


**Supplementary Fig. 10**
